# Single-Cell Indel Detection Enhances Genetic Ancestry and Cellular Lineage Analysis

**DOI:** 10.64898/2026.09.02.748725

**Authors:** Ziyi Wang, Ken Chen, Jinzhuang Dou

**Affiliations:** Department of Bioinformatics and Computational Biology, Division of Basic Science Research, The University of Texas MD Anderson Cancer Center; Graduate Program in Quantitative and Computational Biosciences, Baylor College of Medicine; Department of Biomedical Informatics and Data Science, The University of Alabama at Birmingham

## Abstract

Small insertions and deletions (Indels) provide critical information for cancer genomics and clonal evolution, yet their detection from single-cell sequencing (SCS) such as scRNA-seq and scATAC-seq remains challenging due to sparse coverage, alignment artifacts, and RNA editing. Here, we present Monopogen-Indel, a bioinformatics framework for accurate germline and somatic Indel detection through haplotype-aware variant calling, dynamic template matching in repetitive regions, and cell-population-based allele segregation analysis. We validated germline Indel detection in human retina snRNA-seq with matched bulk whole genome sequencing (WGS). Monopogen-Indel detected 41,000-45,000 germline Indels per sample, with >70% precision and >90% genotyping accuracy. Using 65 heart left ventricle snATAC-seq samples, indel-based global ancestry inference segregated genetic ancestry comparably to SNVs, establishing indels as an independent marker of genetic diversity in SCS. In scRNA-seq from 43,717 cells across four anatomic sites of a patient with high grade serous ovarian cancer (HGSOC), Monopogen-Indel identified ~50,000 germline Indels per sample at 86% WGS-validated precision and an average of 1,040 *de novo* Indels per sample. Approximately 50% of *de novo* Indels reflected biological sources, including variants WGS variants, RNA editing, and cell-type-specific patterns. Somatic SNVs were correctly restricted to CD45^-^ cells, indicating high specificity in delineating somatic from germline variants. In summary, Monopogen-indel is the first framework dedicated to indel detection from SCS and expands the utility of existing single-cell data for population genetics and cancer evolution. The module is integrated into Monopogen repository https://github.com/KChen-lab/Monopogen.

## Introduction

Somatic variants (a.k.a., mutations), including single-nucleotide variants (SNVs), small insertions/deletions (Indels) and structural variants, accumulate in individual cells throughout human lifetime and play critical roles in disease development, particularly in cancer where they could drive tumor initiation, progression and therapeutic resistance (Bley 2020; Pogrebniak and Curtis 2018; Sinkala 2023). Accurate detection of these variants is important for understanding cellular heterogeneity and evolutionary dynamics with the disease. However, conventional bulk sequencing approaches face fundamental limitations in detecting somatic variations, especially those that exist in a limited number of cells and are present at low allelic frequencies (Lawson et al. 2025; Yan et al. 2021). These bulk methods average signals across many cells, obscuring cell-to-cell variation, and limiting the ability to distinguish true somatic variants from technical artifacts and germline polymorphisms. While various techniques have been developed to improve the accuracy of bulk sequencing, such as duplex sequencing (Kennedy et al. 2014), these approaches still operate at the population level and cannot detect mutations in individual cells, which is essential for precisely understanding clonal architecture, including stem or persister cells that drive development and treatment resistance in cancer.

To address these limitations, single-cell DNA sequencing (scDNA-seq) has emerged as a powerful tool for mutation detection at single-cell resolution, directly revealing clonal hierarchies masked in bulk analyses (Miles et al. 2020; Navin et al. 2011). Recently, researchers have also begun leveraging single-cell RNA sequencing (scRNA-seq) and single-cell ATAC sequencing (scATAC-seq) for mutation calling (Alves et al. 2024; Wiens et al. 2024). This shift is driven by the widespread generation of these datasets for profiling gene expression and chromatin accessibility, offering the opportunity to simultaneously link mutations to cellular phenotypes without requiring additional sequencing.

However, existing computational methods for single-cell somatic mutation calling have predominantly focused on SNVs. Tools such as Monovar (Zafar et al. 2016), SComatic (Muyas et al. 2024), and Monopogen (Dou et al. 2024) have demonstrated the feasibility of detecting somatic SNVs from single-cell sequencing data by leveraging information across cells to distinguish true mutations from technical artifacts. Small Indels represent an important class of somatic mutations in cancer (Chen and Guo 2021; Mullaney et al. 2010; Nik-Zainal et al. 2016; Yang et al. 2013). They can not only serve as genetic markers to trace clonal lineages but also have profound functional consequences including frameshift mutations that disrupt tumor suppressors and activating mutations in oncogenes (Alexandrov et al. 2020; Danecek et al. 2021; Fang et al. 2014). Despite their biological significance, Indels have received considerably less attention in SCS-based variant calling, due to additional technical challenges they present, including variable read coverage from SCS, higher error rates in amplification and sequencing (particularly in homopolymer regions), and greater sensitivity to alignment artifacts.

Here, we present Monopogen-Indel, a bioinformatics tool that enables accurate detection of somatic Indels from single-cell sequencing data. Monopogen-Indel’s key innovation is its ability to systematically and efficiently associate observed noisy sequence alignment patterns from individual read to allele segregation patterns expected in a tumor-normal mixed cell population, under the same statistical genetic principle leveraged by Monopogen for somatic SNV calling. Germline SNVs and Indels are present in all cells, whereas somatic Indels are observed in a subset of cells, consistently occurring on the same paternal or maternal alleles across multiple cells due to cellular expansion. It integrates a set of customized bioinformatics filters to reduce Indel calling errors from SCS. By extending Monopogen to include Indel calling, we enable comprehensive single-cell somatic mutation profiling that captures both SNVs and Indels, providing a more complete picture of somatic mutational processes at single-cell resolution.

## Results

### Monopogen-Indel Workflow

Monopogen-Indel focuses on developing new germline and putative somatic Indel calling capability from SCS data, which is not available in the original Monopogen pipeline (Dou et al. 2024). It starts with individual BAM files of SCSs. Variants called from aligned files undergo quality control to remove reads with high alignment mismatches. Putative variants with at least one read supporting the non-reference allele from the pooled pseudobulk (including both SNVs and Indels with length <110 bp) are then collected, and genotype likelihoods are calculated for each site. For any SNVs and Indels present in haplotype reference panels such as 1000 Genomes Phase 3 (1KG3) (Sudmant et al. 2015), linkage disequilibrium (LD) information from the reference panel is collectively used to determine the germline Indels genotypes and phasing information.

Once germline variants are obtained, the pipeline proceeds to somatic Indels variant calling. The pipeline first assigns variant-supporting reads back to individual cells to create a variant-by-cell matrix. To scan the Indel reads in each cell efficiently, we propose the Concurrent Two-Pointer Motif Scan (CTPMScan) strategy. Specifically, we generated a template sequence database spanning the upstream and downstream regions of each candidate variant, including both wild-type and mutant templates, and then sorted the aligned reads by genomic position. We scanned each read to determine whether it matched the wild-type or mutant template based on local positional alignment and subsequently assigned each read’s genotype to its corresponding cell using its barcode. The template sequence was designed to cover 3 bp upstream and downstream of each SNV. To accommodate Indels, which typically occur in repetitive regions and with variable length, we dynamically extend beyond the fixed three-nucleotide window until the flanking repeat sequences terminate (**Method**, **Figure 1**). We then use a support vector machine (SVM) to differentiate true somatic mutations from sequencing errors based on the sequencing alignment features. Finally, we use the cell population pattern to distinguish non-1KG3 germline variants from true somatic variants. This model quantifies whether neighboring Indels co-segregate across the entire population or only within a subpopulation of cells, reflecting their colocalization on a DNA haplotype or RNA transcript.

**Figure 1.**
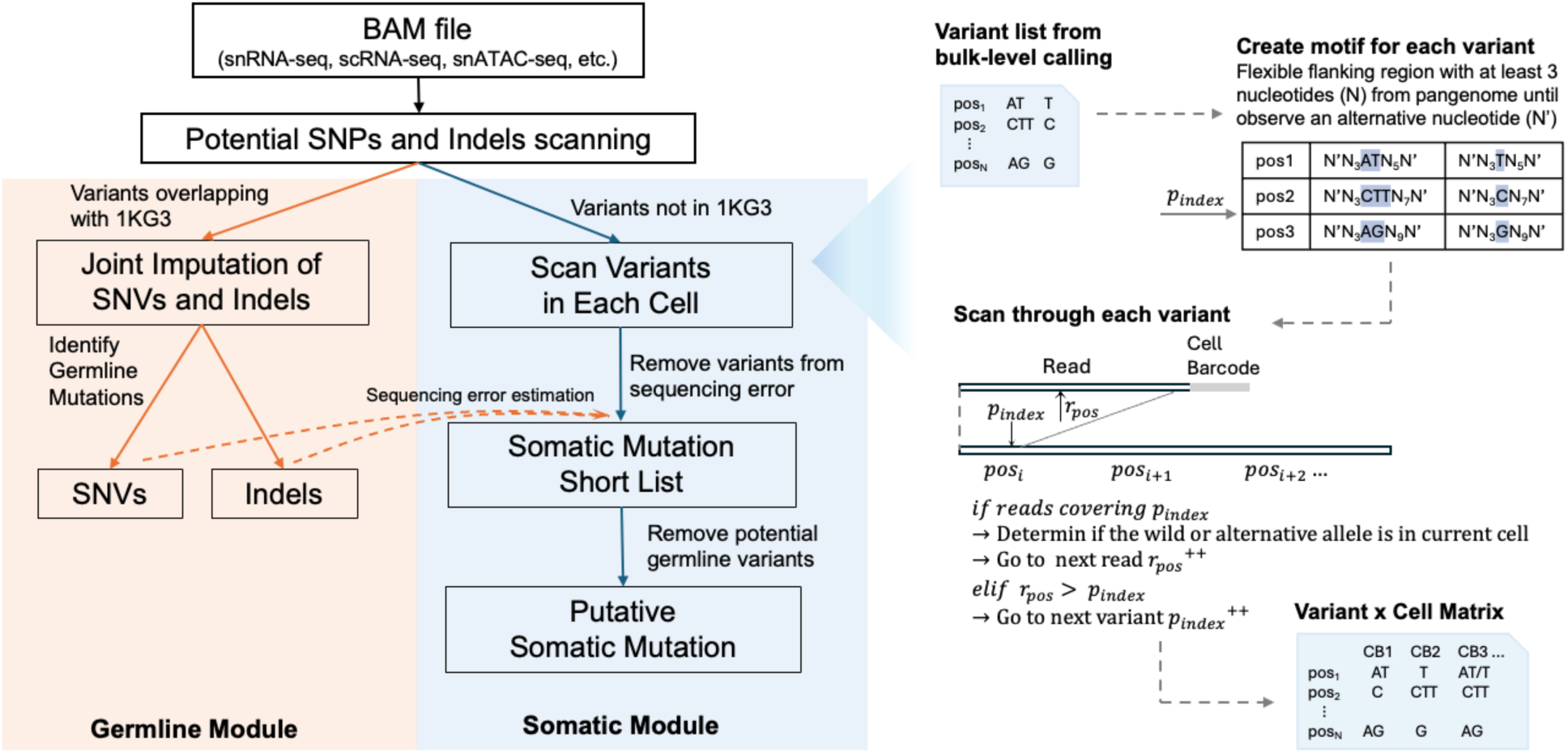
Overview of the Monopogen-indel workflow. The left panel illustrates the germline indel detection and putative somatic indel detection module. The right panel highlights the Concurrent Two-Pointer Motif Scan (CTPMScan) strategy used to generate the cell–variant matrix.

### Validation of Germline Indel Detection Accuracy

To validate Indel calling performance, we used two single-cell RNA sequencing datasets with matched WGS data: (1) snRNA-seq data from four retina tissue samples, and (2) scRNA-seq multi-tissue data from the MSK-SPECTRUM study of 8 high-grade serous ovarian cancer (Vázquez-García et al. 2022). For retina snRNA-seq data, the Monopogen-Indel detected on average of 42.6k (41,016 – 45,734) germline Indels per sample. When defining the gold standard as having minimum 4 supporting reads in the WGS, Monopogen-Indel detected an average of 31,145 Indels overlapping with that gold standard, yielding an average recall rate of 3.93%, and precision of 73.0% in callable regions (**Table 1**). The Indels range from –308 bp to +949 bp, with 95% interval from –15bp to +13bp. The recall rate is lower than that of SNV with (~20%) in the same sample (Dou et al. 2024). For Indels overlapped with the WGS panel, genotyping accuracy exceeded 90% for all four samples when validated against WGS data. Most genotyping errors involved misclassification between homozygous and heterozygous calls (1/1 to 0/1 or 0/1 to 1/1).

**Table 1.** Total germline variants detected from retina snRNA-seq.

| Sample ID | No. of Indel in WGS | No. of Indel in snRNA seq | No. of Indels overlapping | 0/1->1/1 | 1/1->0/1 | Recall | Precision | Genotyping accuracy |
| --- | --- | --- | --- | --- | --- | --- | --- | --- |
| 19D013 | 818 167 | 45 734 | 33 487 | 825 | 865 | 4.1% | 73.2% | 92.0% |
| 19D014 | 780 458 | 41 823 | 29 972 | 764 | 842 | 3.8% | 71.7% | 91.6% |
| 19D015 | 803 820 | 41 016 | 30 563 | 625 | 777 | 3.8% | 74.5% | 92.5% |
| 19D016 | 759 730 | 42 033 | 30 557 | 626 | 920 | 4.0% | 72.7% | 92.0% |

In the MSK spectrum data, we benchmarked performance on a randomly selected patient sample collected from 4 anatomic sites, with cells stratified by CD45 status as in the original study (Vázquez-García et al. 2022). The pipeline detected on average 56,465 (52,080-60,248) germline Indels per dataset, with a calling precision around 87% (**Table 2**). The pipeline detected on average 27 (16-39) somatic Indels per dataset, with calling precision around 3% (**Table 3**). We also called germline SNVs using Monopogen default settings on this dataset. Again, comparison between variant types revealed that germline Indel calling achieved lower precision than SNV calling (87% vs. 97%), and lower recall (11.31% vs. 28.82% on average). We observed no significant differences in the number of variants called across different sampling sites or cell types.

**Table 2.**
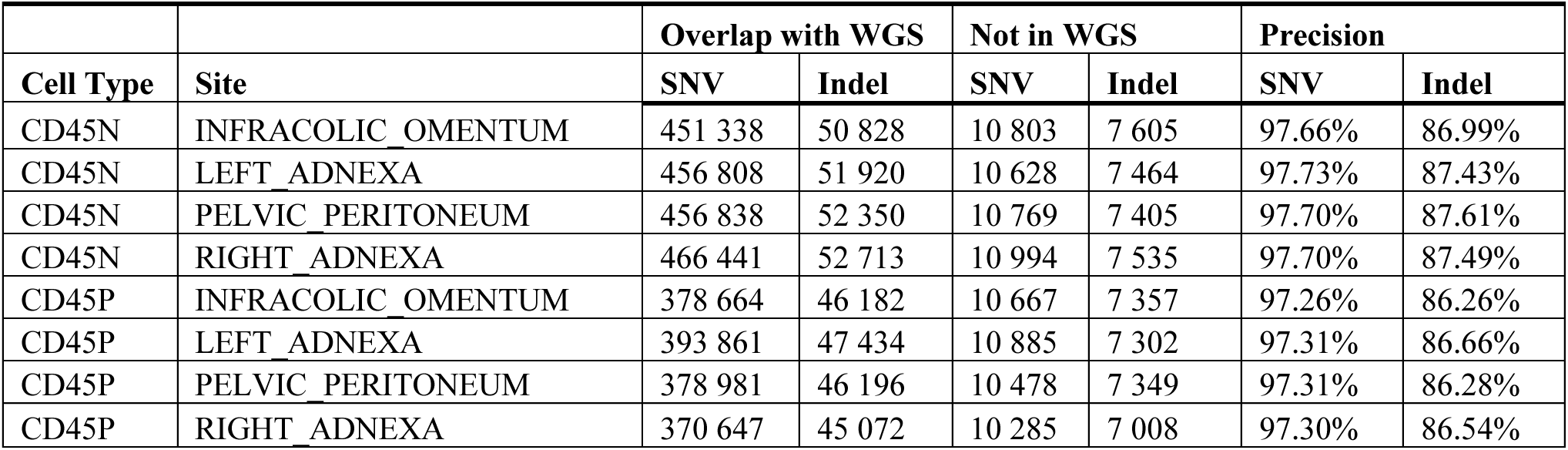
Total germline variants detected from MSK Spectrum HGSOC samples.

**Table 3.** Total somatic variants detected from MSK Spectrum HGSOC samples.

|  |  | Overlap with WGS Germline |  | Overlap with WGS Tumor |  | Not in WGS |  | Precision |  |
| --- | --- | --- | --- | --- | --- | --- | --- | --- | --- |
| Cell Type | Site | SNV | Indel | SNV | Indel | SNV | Indel | SNV | Indel |
| CD45N | INFRACOLIC OMENTUM | 533 | 16 | 24 | 0 | 7 520 | 668 | 6.90% | 2.34% |
| CD45N | LEFT ADNEXA | 570 | 24 | 6 | 0 | 7 043 | 924 | 7.56% | 2.53% |
| CD45N | PELVIC PERITONEUM | 576 | 20 | 13 | 0 | 7 160 | 995 | 7.60% | 1.97% |
| CD45N | RIGHT ADNEXA | 568 | 24 | 14 | 0 | 7 361 | 843 | 7.33% | 2.77% |
| CD45P | INFRACOLIC OMENTUM | 612 | 32 | 0 | 0 | 7 918 | 1 175 | 7.17% | 2.65% |
| CD45P | LEFT ADNEXA | 657 | 31 | 0 | 0 | 7 131 | 1 208 | 8.44% | 2.50% |
| CD45P | PELVIC PERITONEUM | 641 | 39 | 0 | 0 | 7 152 | 1 199 | 8.23% | 3.15% |
| CD45P | RIGHT ADNEXA | 568 | 31 | 0 | 0 | 6 811 | 1 087 | 7.70 | 2.77% |

### Global Ancestry Inference with germline Indels from SCS Data

To evaluate whether germline Indels provide comparable power to SNVs for genetic ancestry inference at the population level, we perform the germline SNVs and Indels detection on 65 donors from the snATAC-seq data of the ENCODE human heart left ventricle dataset (Eraslan et al. 2022). With the obtained germline SNVs, we projected donor genotypes onto a global ancestry reference panel from the Human Genome Diversity Project (HGDP) (J. Z. Li et al. 2008), which includes populations from East Asia, America, the Middle East, Europe, Oceania, Africa, and Central/South Asia. Such projection enables us to establish ancestry assignments using SNP-based PCA as the ground truth. We then performed PCA using the 1,249,093 detected germline Indels and compared the resulting population structure to the SNP-based assignments (**Figure 2**). Although the number of Indels are lower, the Indel-based ancestry inference recapitulated the clustering patterns observed with SNP-based inference, demonstrating that Indels provide independent, discriminatory power for ancestry mapping in this dataset, which could lead to deeper delineation of the population substructure.

**Figure 2.**
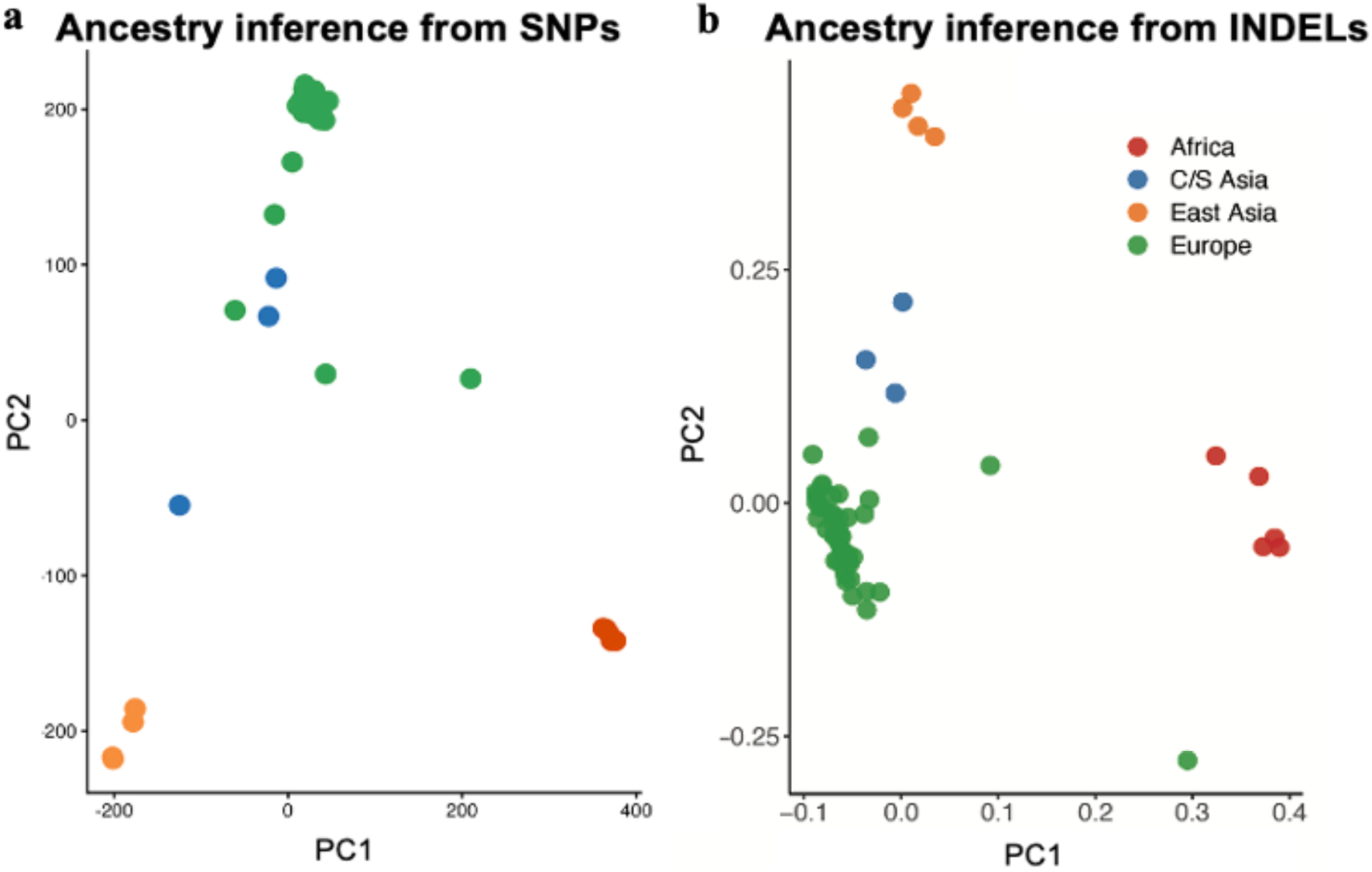
a. Projection of ENCODE heart left ventricle snATAC-seq samples onto the HGDP reference panel based on SNP genotypes inferred using Monopogen. Samples are colored by ancestry classification determined in the projection space, using HGDP as the reference. b. PCA of indel genotypes inferred with Monopogen-indel. Samples are colored according to ancestry labels assigned from SNP-based classification.

### Somatic Indel Detection Performance from SCS Data

To evaluate the detection power of Monopogen–Indels on detecting somatic Indels, we examined 43,717 cells from a patient with high-grade serous ovarian cancer (HGSOC). These cells were collected from 4 anatomic sampling sites, sorted into CD45^+^ or CD45^-^ populations. Using matched WGS data from both the primary tumor site and Peripheral Blood Mononuclear Cells (PBMCs), we identified 1.45M germline SNVs and 433k germline Indels, as well as 2,420 tumor-specific somatic SNVs and 300 tumor-specific somatic Indels (**Methods**; **Figure 3**)

**Figure 3.**
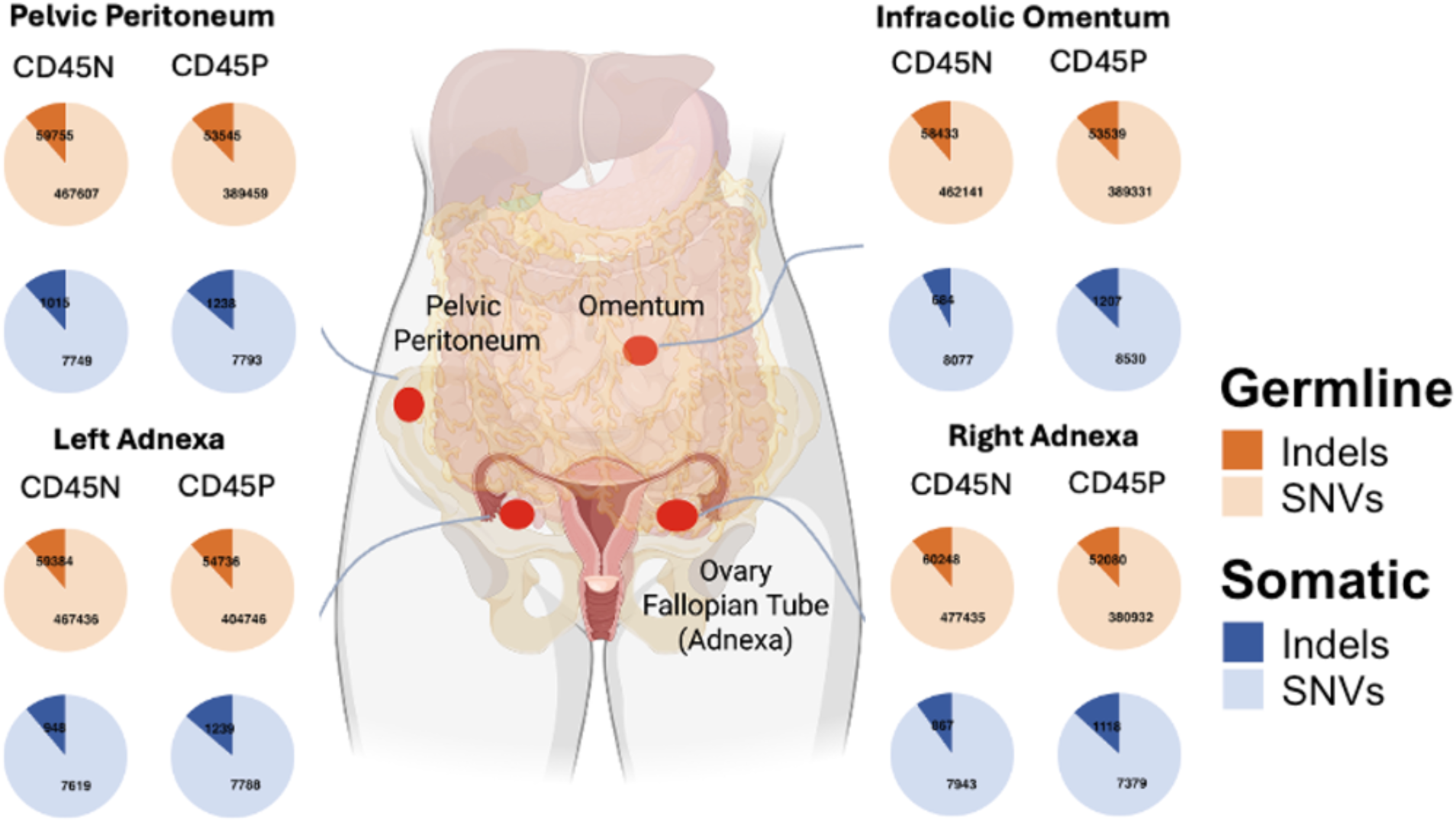
Variant calling summary from 4 sites of the same donor with high-grade serous ovarian cancer samples in MSK-SPECTRUM study.

We applied Monopogen-Indel to identify somatic SNVs and Indels from each anatomic sampling site from SCSs and compared them to variants detected from the WGS data of the same patient samples (**Table 3**). On average, we identified 1,040 Indels per sample (range: 684-1,239, size range: −14 bp - +9 bp), from SCSs. There was an average of 27 Indels (range: 16-39) overlapping WGS germline variants, while none overlapped with WGS somatic Indels, due likely to the low sensitivity of Indel detection in SCS and in WGS. To evaluate the specificity of somatic variant calling, we examined the somatic SNV calls generated from the same pipeline. All SNVs overlapping with WGS tumor somatic mutations originated from CD45^-^ cells, confirming that Monopogen-Indel can correctly identify tumor cells and distinguish them from CD45^+^ immune cells. Similar to germline detection, more somatic SNVs were identified than somatic Indels, and the precision for SNV detection (7.62%) was relatively higher than for Indels (2.59%).

### Characterization of *de novo* Somatic Variants

To identify the sources of the 58,096 *de novo* SNVs and 8,099 *de novo* Indels and determine whether they represented sequencing artifacts or true mutations missed by the WGS pipeline, we performed a systematic characterization. We first examined the number of sampling sites sharing each variant (**Figure 4a-b**). The analysis revealed a significant number of variants (65.79% germline SNVs, 49.86% germline Indels, 32.76% somatic SNVs, 15.93% somatic Indels) occurring in more than 2 samples, suggesting that a large fraction of the *de novo* variants are bona fide variants missed by the WGS.

**Figure 4.**
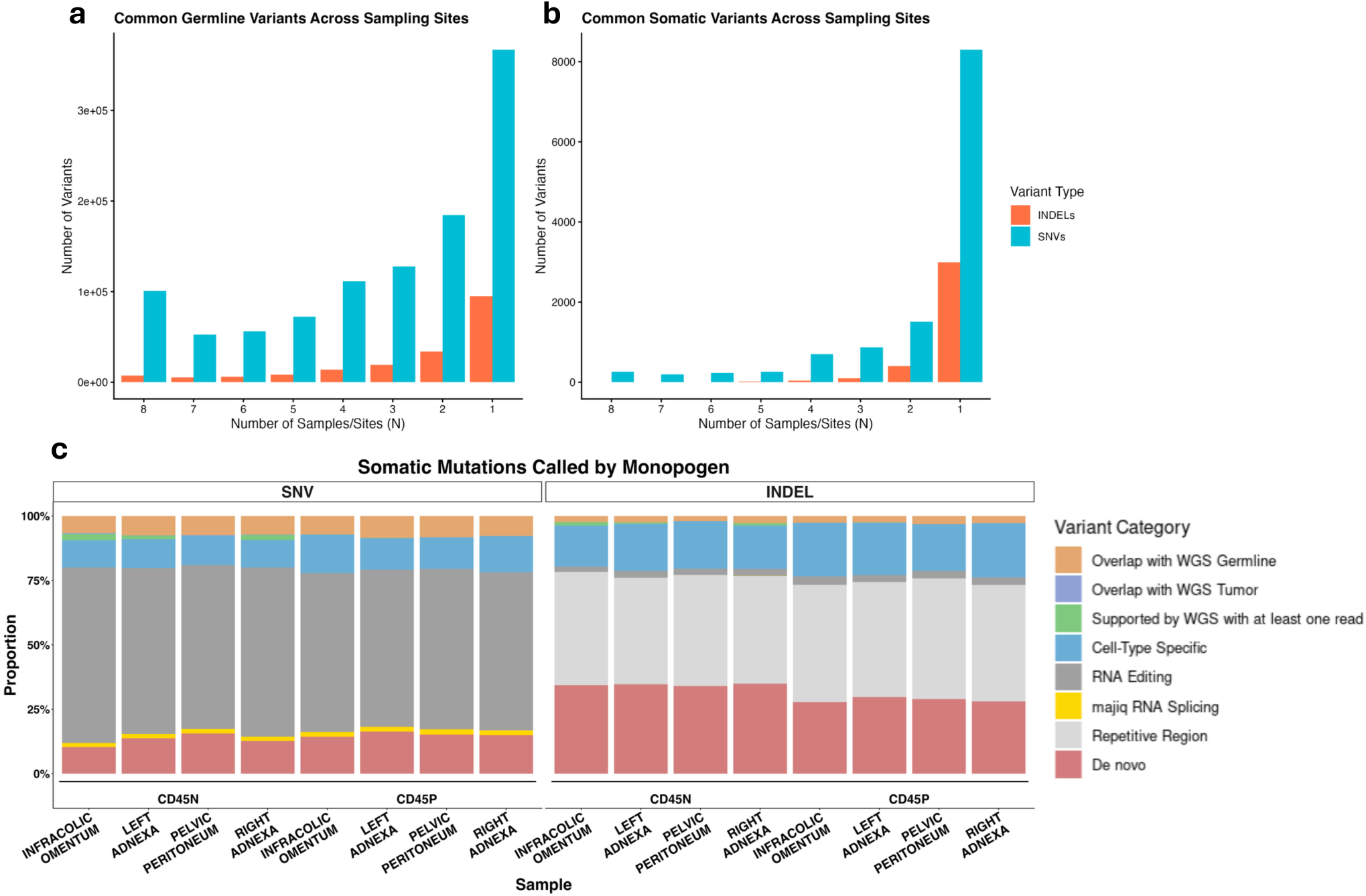
a. Common germline variants shared among number of sampling sites. b. Common somatic variants shared among number of sampling sites. c. Somatic variants called by Monopogen categorized by their source.

We then systematically categorized these *de novo* variants based on multiple lines of evidence (**Figure 4c**). First, we examined WGS pileup files to determine whether reads supporting these variants were present but filtered out by the WGS pipeline. On average, each sampling site has 590 SNVs and 27 Indels overlapping with WGS germline variants, and 7 SNVs and 0 Indels overlapping with WGS tumor. Then, we checked whether the somatic variants were overlapping with known RNA editing sites (Kiran et al. 2013; Picardi et al. 2017; Tan et al. 2017) and *de novo* RNA editing sites based on local splicing variations (Aicher et al. 2024). On average, each site has 4,992 SNVs and 29 Indels in the RNA editing database. For the *de novo* editing sites, there are 103 SNVs and 1 Indel on average for each site. Third, because Indels frequently occur in repetitive regions, we assessed overlap with regions identified by RepeatMasker (Bao and Eddy 2002). All the Indels identified falls in the repetitive regions, with an average of 460 Indels per sequencing site. Finally, we evaluated whether de novo mutations exhibited cell-type-specific patterns (**Methods, Figure 5**). As shown in Fig. 5, We detected Indels specifically in tumor cells but not in lymphocytes, similar to other SNV alleles, supporting their somatic nature. Interestingly, some Indels were detected in both tumor and fibroblast cells, indicating potential cellular plasticity. Through such categorization, we could attribute approximately 55% of *de novo* Indels, with the remainder likely representing sequencing errors or resulting from unknown factors. Notably, the categorization was more successful for SNVs, explaining approximately 79% of *de novo* variants (**Supplementary Table 1**). This difference also suggests that Indel detection remains more challenging and prone to artifacts than SNV detection in single-cell data.

**Figure 5.**
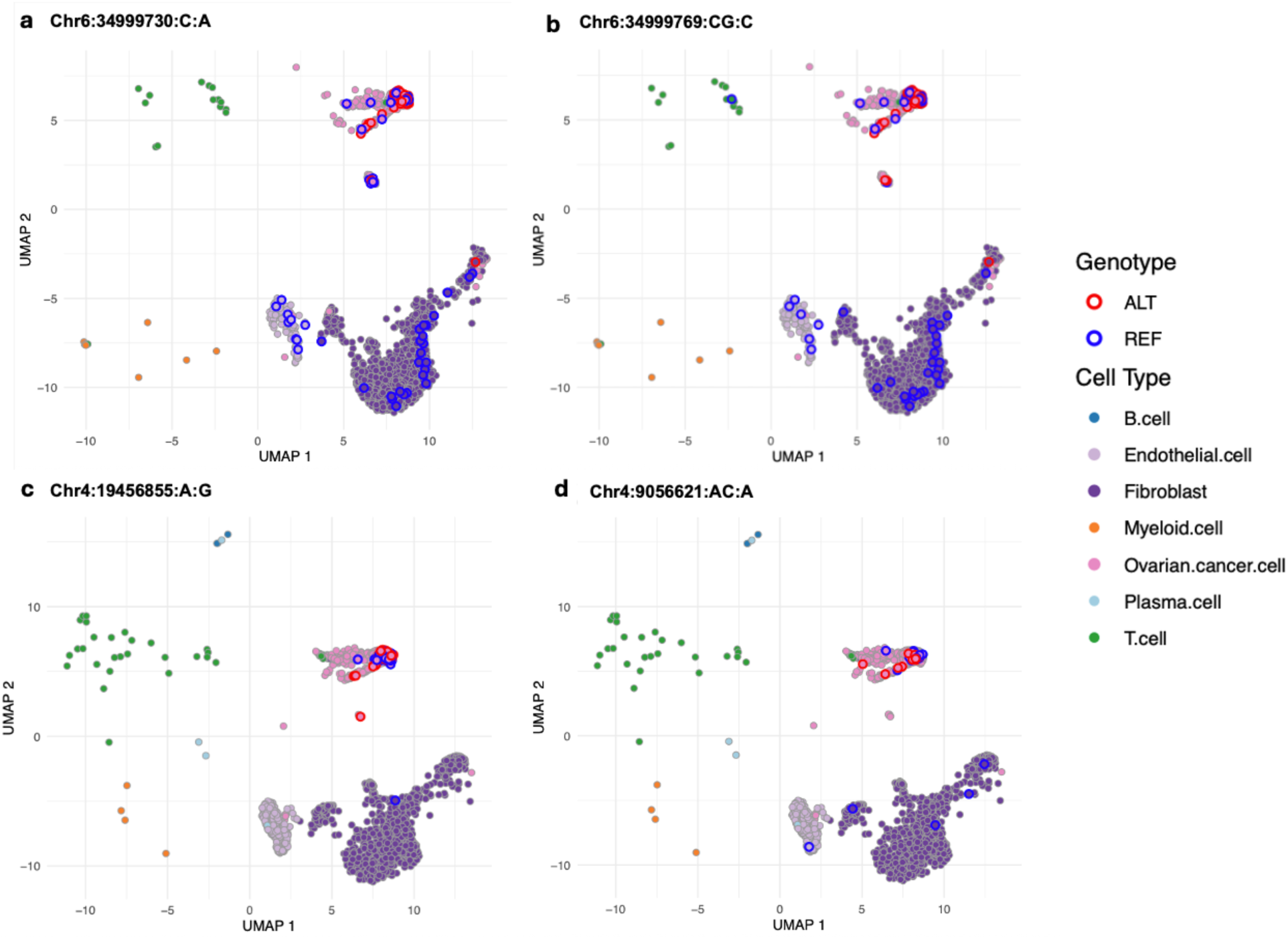
Representative examples of somatic indel distributions across cell populations. (a) Somatic SNV Chr6:34999730:C: A. (b) Somatic Indel chr6:34999769:CG: C. (c) Somatic SNV Chr4:19456855: A:G. (d) Somatic Indel chr4:9056621:AC: A. The right panel presents representative somatic Indels exhibiting distribution patterns comparable to those observed for the somatic SNVs.

## Discussion

In this study, we developed Monopogen-indel to detect Indel variants from SCS data, enabling comprehensive characterization of both SNVs and Indels at single-cell resolution. Our analysis demonstrated that Monopogen-Indel can successfully identify Indel variants while maintaining cell level resolution. We show that Indels provide information complementary to SNVs for resolving fine-scale population structure for single cell population genetics and distinguishing closely related cell lineages within heterogeneous samples. These results establish a framework for joint SNV and Indel analysis in SCS data, with broad applications for cancer clonal evolution studies, developmental biology, and population genetics at single-cell resolution.

Short Indels, particularly those less than 5 base pairs, are inherently prone to multiple error sources affecting both germline and somatic variant detection (H. Li 2014). These variants are susceptible to PCR slippage in homopolymer region (Fang et al. 2014; Fungtammasan et al. 2015), sequencing errors (Stoler and Nekrutenko 2021), and alignment artifacts (Albers et al. 2011). In scRNA-seq, where per-cell coverage is limited and allelic dropout is common (Wiens et al. 2024), distinguishing true short Indels from technical noise is particularly challenging (Sun et al. 2017). Similarly, in scATAC-seq, the per-locus fragments are sparce, and Tn5 transposition might also introduce additional sequencing biases (Hu et al. 2022). To mitigate these issues, we filtered variants against the RepeatMasker database and RNA editing sites to exclude potential technical artifacts, but some residual errors likely remain, particularly in repetitive genomic contexts.

In our model, tumor purity could also affect somatic variant detection accuracy through LD shifts. As tumor purity decreases, somatic variants result in less sequencing of reads, becoming hard to detect. Such effects likely impact scRNA-seq data more severely than scATAC-seq data, as allele-specific and cell-type specific gene expression can magnify LD distortions. Additionally, while we validated that our approach captures cell type-specific regulatory effects in both normal and disease states, germline variants with allele-specific expression can be mistaken for somatic variants when one allele is preferentially expressed. This is particularly problematic in tumors with low purity or exhibit allelic imbalance and represents an important consideration for future applications of single-cell Indel detection in heterogeneous tumor samples.

Determining whether detected *de novo* mutations represent true biological events or sequencing artifacts remains challenging, especially for Indels which have higher error rates than SNVs. We acknowledge that some detected Indels, particularly those with low allelic frequencies in low-complexity regions, may represent technical artifacts. In the future, we plan to add additional reference panels such as TOPMed (Taliun et al. 2021), All of Us (All of Us Research Program et al. 2019), and UK Biobank (Sudlow et al. 2015) to account for ethnic disparity. Additionally, we plan to incorporate other validation methods such as targeted single-cell DNA sequencing or long-read technologies that would help distinguish true mutations from sequencing errors.

Despite these limitations, our extension of Monopogen to include Indel detection represents an important advance for single-cell genomics, enabling more comprehensive mutation profiling that captures the full spectrum of variants driving cellular phenotypes and disease progression.

## Acknowledgements

This project was supported from Chan Zuckerberg Initiative Donor Advised Fund (CZI DAF) [DAF2024-345892 to R. C and K. C]

## Data Availability Statement

The datasets used in this study are publicly available. The snRNA-seq and snATAC-seq profiles from the human heart left ventricle tissues of 65 donors were downloaded from ENCODE study60 at https://www.encodeproject.org/matrix/?type=Experiment&assay_title=snATAC-seq&assay_title=scRNA-seq&biosample_ontology.term_name=heart+left+ventricle. The 1KG3 genotypes were from 1000 genome project and downloaded from https://ftp.1000genomes.ebi.ac.uk/vol1/ftp/data_collections/1000G_2504_high_coverage/working/20201028_3202_phased/. The HGDP panel62 genotypes were downloaded from http://csg.sph.umich.edu/chaolong/LASER/HGDP-938-632958.tar.gz. The four retina single-cell samples could be downloaded from https://data.humancellatlas.org/explore/projects/f0f89c14-7460-4bab-9d42-22228a91f185. The MSK-SPECTRUM data was downloaded from https://www.synapse.org/Synapse:syn25569736/wiki/612269. The restricted sequencing files are requested from dbGaP, study accession: phs002857.v3.p1.

The implementation code is publicly in https://github.com/KChen-lab/Monopogen.

## Method

### Monopogen-Indel workflow

#### Preprocessing and Quality Control

Data processing began with BAM files from individual single-cell samples, which were converted to variant call format through a coordinated Samtools and BCFtools workflow (Danecek et al. 2021; H. Li et al. 2009). We implemented stringent quality thresholds to ensure high-confidence variant detection: sequencing reads failing to meet a minimum mapping quality of 20 were discarded, and individual base calls with quality scores below 20 were excluded from pileup generation. The initial variant call set then underwent systematic filtering to remove positions with ambiguous or non-standard reference base calls, including symbolic allele representations that could confound downstream analysis. Complex variant sites harboring multiple alternative alleles were removed due to their complexity in downstream genotype assignments. All resulting variant records were subsequently left-aligned and normalized relative to the reference genome assembly.

#### Germline Variant Detection

Quality-controlled VCF files were processed through Monopogen’s germline detection module, which utilized haplotype data from the 1000 Genomes Project Phase 3 (1KG3) reference panel to infer genotype. By integrating genotype likelihoods with population-level LD patterns, the pipeline phased single-cell alignments and assigned corresponding genotypes.

#### Somatic Variant Detection

To identify somatic mutations, we focus on candidate loci that is not overlapped in 1KG3 reference panel. We applied population-level linkage patterns as an additional validation layer, under the assumption that germline variants maintain characteristic co-occurrence patterns with neighboring polymorphisms, whereas true somatic mutations lack these inherited correlations. This filtering effectively distinguished residual germline variants absent from reference databases from genuine somatic alterations.

After establishing the potential somatic variant list, we constructed single-cell resolution genotype matrices by tracing variant-supporting sequencing reads back to their cells of origin. To efficiently scan reads in each cell, we propose the CTPMScan strategy. Specifically, we generated a template sequence database spanning the upstream and downstream regions of each candidate variant, including both wild-type and mutant templates, and then sorted the aligned reads by genomic position. We scanned each read to determine whether it matched the wild-type or mutant template. To avoid exhaustive pairwise scanning between reads and the candidate variant list, we introduced a positional index: reads are only scanned starting from the current index position, and once a match is found, the search is terminated, and the index is updated accordingly. Based on this process, each read’s genotype is assigned to its corresponding cell using its barcode. The template sequence was designed to cover 3 bp upstream and downstream of each SNV. To accommodate Indels, which typically occur in repetitive regions and with variable length, we developed an adaptive template strategy. Instead of using a fixed flanking window as in the original Monopogen pipeline, our current templates expand iteratively until reaching non-repetitive sequence boundaries, ensuring robust variant detection regardless of Indel complexity.

The final classification stage integrates reference-panel-informed error modeling with machine learning discrimination. A support vector machine classifier, trained on error rates derived from phased germline genotypes, separates technical artifacts from biological mutations. Subsequently, cell population informed post-processing resolves ambiguous cases where novel germline polymorphisms (absent from 1KG3) might otherwise be misclassified as somatic events. The Indel module has been implemented and integrated into the Monopogen repository https://github.com/KChen-lab/Monopogen.

### Germline Variant Calling Evaluation

We performed a germline variant calling evaluation on 12 samples, comprising 4 retina samples and 8 high-grade serous ovarian carcinoma (HGSOC) samples.

#### Performance Metrics

For performance comparison, detected mutations were classified into three categories: (1) true positives (TP): variants identified in both the single-cell dataset and the matched WGS data; (2) false positives (FP): variants called in the single-cell dataset but absent from the WGS data; and (3) false negatives (FN): variants present in the WGS data within genomic regions covered by the single-cell assay but not detected by the single-cell method. Precision and recall were calculated for each method using standard formulas: precision = TP / (TP + FP) and recall = TP / (TP + FN).

#### Genotyping Accuracy Assessment

To evaluate Monopogen’s genotyping accuracy, we analyzed 4 retina single-cell samples with matched WGS data. For each sample, we extracted bi-allelic loci containing at least one alternative allele from both Monopogen and WGS call sets. Performance was assessed using multiple metrics: recall quantified the proportion of WGS-called variants successfully detected by Monopogen, precision measured the proportion of Monopogen-called variants validated by WGS, and genotyping accuracy represented the fraction of overlapping variants with concordant genotype assignments between both methods.

### Global Ancestry Analysis

To determine the global ancestry of single-cell sequencing samples, we obtained genotype data from the Human Genome Diversity Project (HGDP) (Bergstrom et al. 2020), comprising 938 individuals representing 53 worldwide populations and 632 958 single nucleotide variants (SNVs) with minor allele frequency (MAF) > 1%. Each sample was projected onto the HGDP reference panel using the LASER software (Trace module)(Liu et al. 2014).

We analyzed 65 donor samples from the ENCODE human left ventricle dataset. Germline SNPs identified by Monopogen were first used to classify samples into broad ancestry categories: African, Central/South Asian, East Asian, and European. Subsequently, principal component analysis (PCA) was performed on germline Indels, with samples colored according to their SNP-derived ancestry classifications to visualize population structure and validate ancestry assignments.

### Somatic Variant Calling Evaluation

To establish a high-confidence somatic variant reference set, we obtained matched WGS FASTQ files from primary tumor tissue and peripheral blood mononuclear cells (PBMCs) of Donor 115 from the MSK-SPECTRUM dataset via dbGaP, study accession: phs002857.v3.p1 (Vázquez-García et al. 2022). Raw sequencing reads were aligned to the human reference genome GRCh38 using BWA-MEM 0.7.18-r1243-dirty (H. Li and Durbin 2009), followed by quality control with GATK v4.6.1.0 (Van der Auwera et al. 2020), including PCR duplicate marking and base quality score recalibration. Ground truth classification was determined through bidirectional Mutect2 (Cibulskis et al. 2013) comparison: tumor versus PBMC and PBMC versus tumor. Variants identified as somatic in either direction were designated as tissue-specific somatic variants, while those consistently absent from somatic calls in both comparisons were classified as germline variants.

Numeric performance assessment for somatic variants was conducted using methods analogous to those applied for germline variants. Additionally, we evaluated the precision of somatic variant calls by comparing WGS-derived tumor somatic variants against WGS germline variants to assess the specificity of somatic detection.

#### *De Novo* Variant Characterization

To comprehensively characterize putative de novo variants, we applied multiple filtering and annotation strategies. Variants were systematically annotated and categorized based on known RNA editing sites (Kiran et al. 2013; Picardi et al. 2017; Tan et al. 2017), majiq3 de novo RNA splicing detection (Aicher et al. 2024), repetitive genomic regions (Smit 2013–2015), and evidence from WGS read pileups. Specifically, we examined raw pileup files from matched WGS data to assess the presence of supporting reads for each candidate variant.

To evaluate potential cell-type specificity of de novo variants, we incorporated cell type annotations from the original publication. We then performed Fisher’s exact test to identify statistically significant associations between variant presence and specific cell types, enabling the detection of cell-type-enriched mutational patterns.

---

**Supplementary Table 1.**
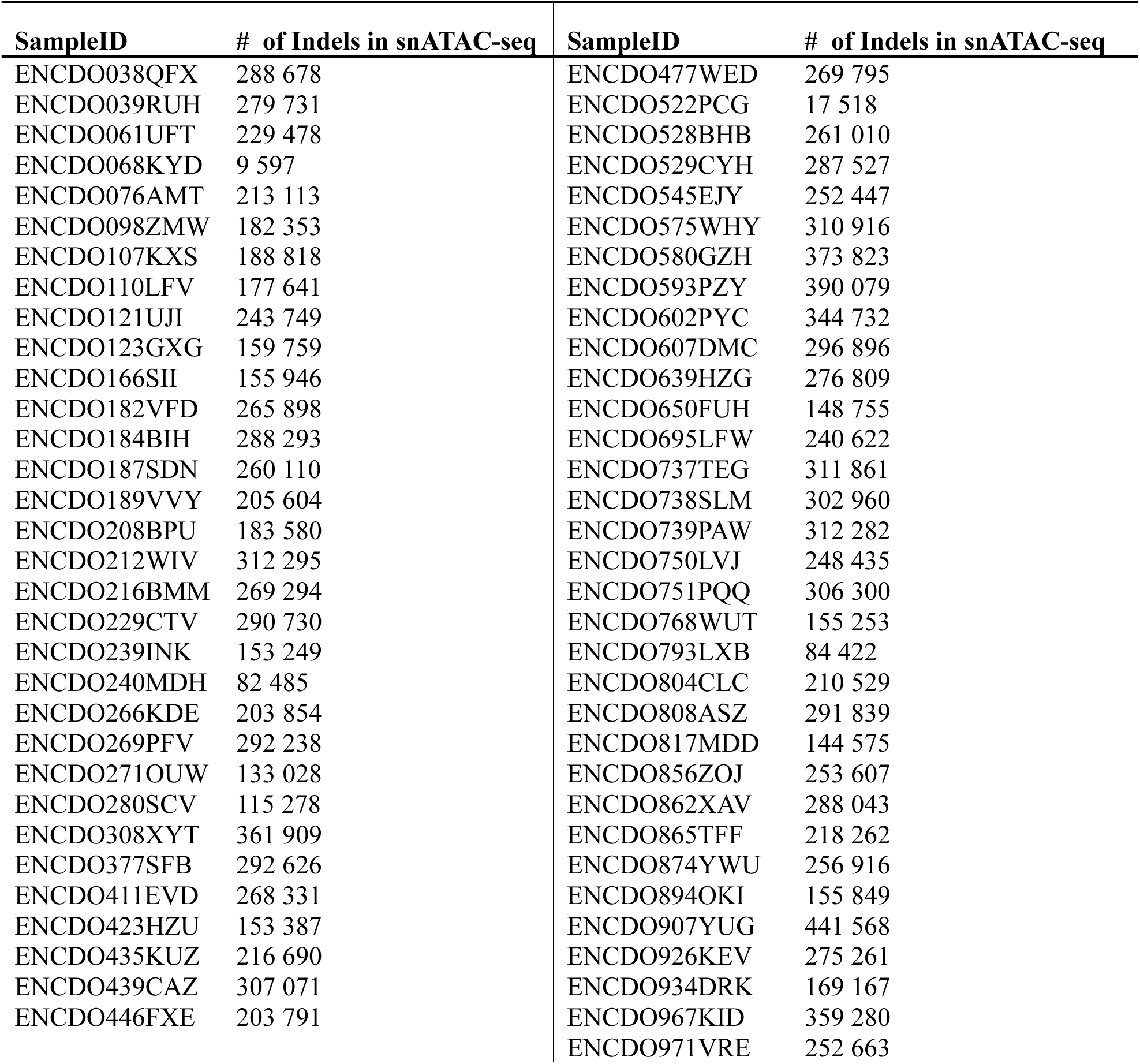
Summary of Indel calling from heart left ventricle samples.

**Supplementary Figure 1.**
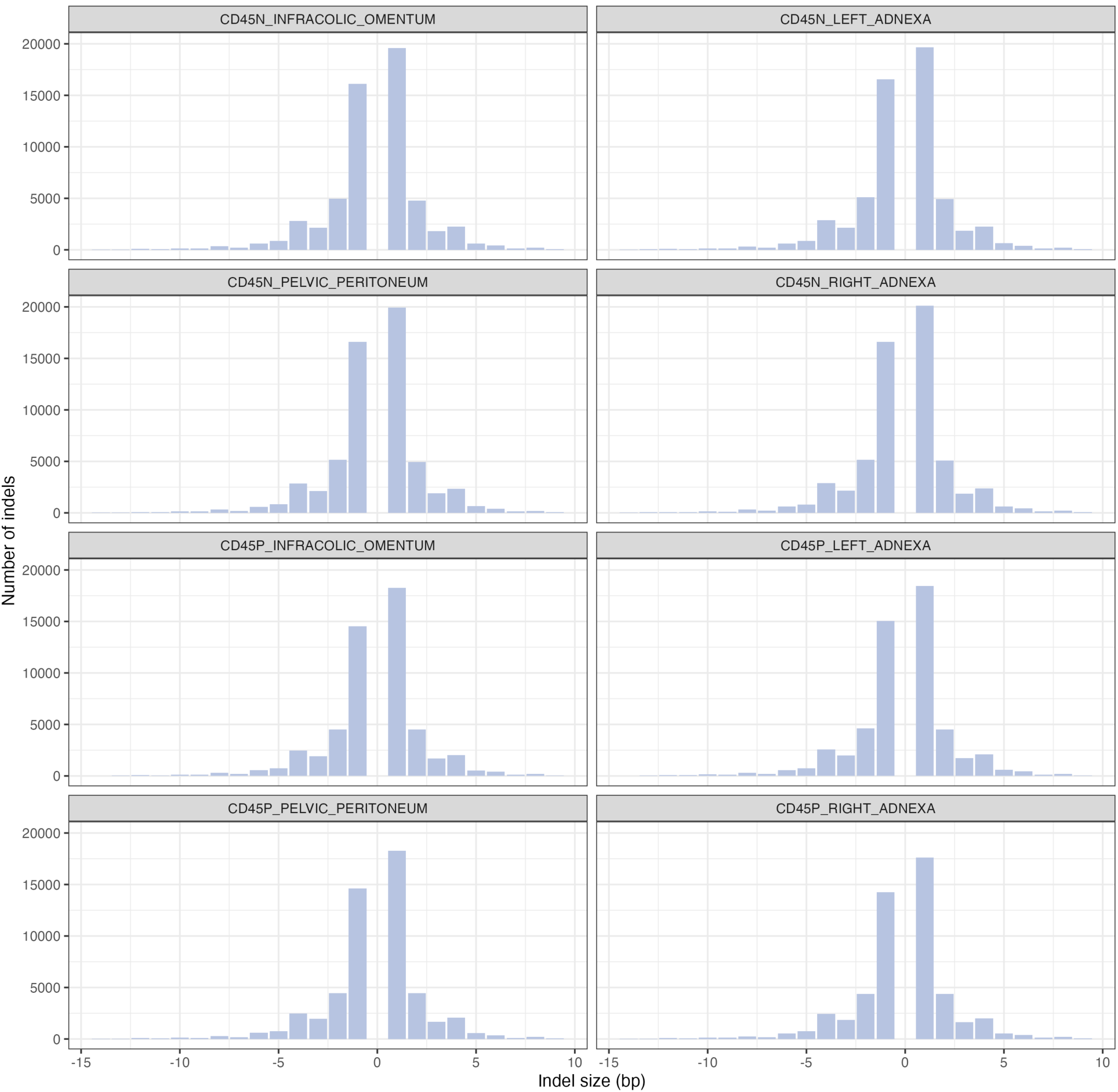
Distribution of insertion and deletion sizes of germline indels on HGSOC. Positive values represent insertion lengths, whereas negative values represent deletion lengths.

**Supplementary Figure 2.**
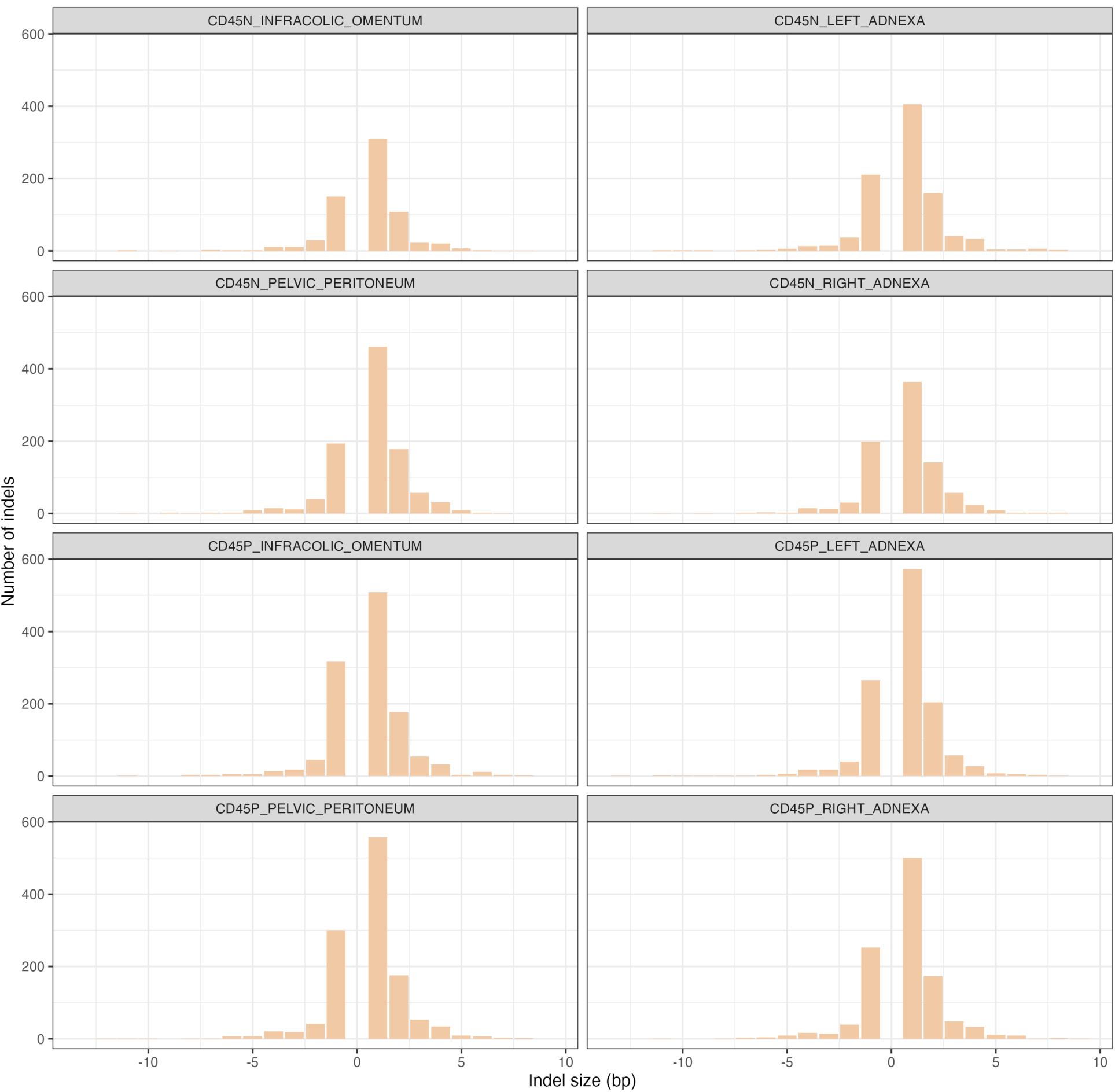
Distribution of insertion and deletion sizes of somatic indels on HGSOC. Positive values represent insertion lengths, whereas negative values represent deletion lengths.

**Supplementary Figure 3.**
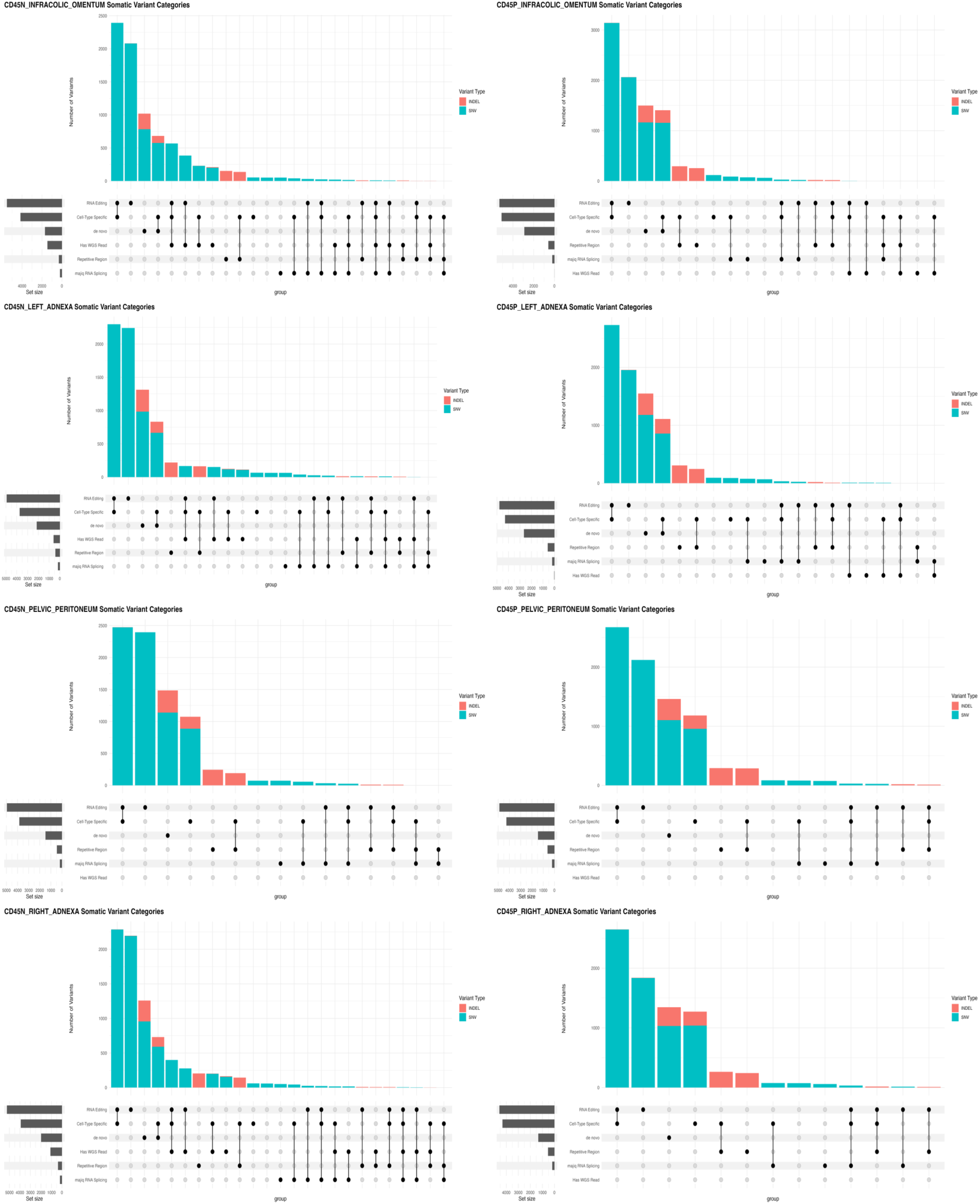
Somatic variants called at each sampling site, categorized by type.

## Reference

Aicher, Joseph K, et al. (2024), ‘MAJIQ V3 offers improvements in accuracy, performance, and usability for splicing analysis from RNA sequencing’, bioRxiv, 2024.07.02.601792.

Albers, C. A., et al. (2011), ‘Dindel: Accurate indel calls from short-read data’, Genome Research, 21 (6), 961–73.

Alexandrov, L. B., et al. (2020), ‘The repertoire of mutational signatures in human cancer’, Nature, 578 (7793), 94–101.

All of Us Research Program, Investigators, et al. (2019), ‘The “All of Us” Research Program’, N Engl J Med, 381 (7), 668–76.

Alves, Joao M, Tomás, Laura, and Posada, David (2024), ‘Unraveling the phylogenetic signal of gene expression from single-cell RNA-seq data’, bioRxiv, 2024.04.17.589871.

Bao, Z. R. and Eddy, S. R. (2002), ‘Automated de novo identification of repeat sequence families in sequenced genomes’, Genome Research, 12 (8), 1269–76.

Bergstrom, A., et al. (2020), ‘Insights into human genetic variation and population history from 929 diverse genomes’, Science, 367 (6484).

Bley, N. (2020), ‘TUMOR EVOLUTION Finding the mutations that drive resistance’, Elife, 9.

Chen, J. and Guo, J. T. (2021), ‘Structural and functional analysis of somatic coding and UTR indels in breast and lung cancer genomes’, Scientific Reports, 11 (1).

Cibulskis, K., et al. (2013), ‘Sensitive detection of somatic point mutations in impure and heterogeneous cancer samples’, Nature Biotechnology, 31 (3), 213–19.

Danecek, P., et al. (2021), ‘Twelve years of SAMtools and BCFtools’, Gigascience, 10 (2).

Dou, J. Z., et al. (2024), ‘Single-nucleotide variant calling in single-cell sequencing data with Monopogen’, Nature Biotechnology, 42 (5), 803–+.

Eraslan, G., et al. (2022), ‘Single-nucleus cross-tissue molecular reference maps toward understanding disease gene function’, Science, 376 (6594), 712–+.

Fang, H., et al. (2014), ‘Reducing INDEL calling errors in whole genome and exome sequencing data’, Genome Medicine, 6.

Fungtammasan, A., et al. (2015), ‘Accurate typing of short tandem repeats from genome-wide sequencing data and its applications’, Genome Res, 25 (5), 736–49.

Hu, S. G. S., et al. (2022), ‘Intrinsic bias estimation for improved analysis of bulk and single-cell chromatin accessibility profiles using SELMA’, Nature Communications, 13 (1).

Kennedy, S. R., et al. (2014), ‘Detecting ultralow-frequency mutations by Duplex Sequencing’, Nature Protocols, 9 (11), 2586–606.

Kiran, A. M., et al. (2013), ‘Darned in 2013: inclusion of model organisms and linking with Wikipedia’, Nucleic Acids Research, 41 (D1), D258–D61.

Lawson, A. R. J., et al. (2025), ‘Somatic mutation and selection at population scale’, Nature, 647 (8089), 411–20.

Li, H. (2014), ‘Toward better understanding of artifacts in variant calling from high-coverage samples’, Bioinformatics, 30 (20), 2843–51.

Li, H. and Durbin, R. (2009), ‘Fast and accurate short read alignment with Burrows-Wheeler transform’, Bioinformatics, 25 (14), 1754–60.

Li, H., et al. (2009), ‘The Sequence Alignment/Map format and SAMtools’, Bioinformatics, 25 (16), 2078–79.

Li, J. Z., et al. (2008), ‘Worldwide human relationships inferred from genome-wide patterns of variation’, Science, 319 (5866), 1100–04.

Liu, H., et al. (2014), ‘Rare earth elements recycling from waste phosphor by dual hydrochloric acid dissolution’, J Hazard Mater, 272, 96–101.

Miles, L. A., et al. (2020), ‘Single-cell mutation analysis of clonal evolution in myeloid malignancies’, Nature, 587 (7834), 477–82.

Mullaney, J. M., et al. (2010), ‘Small insertions and deletions (INDELs) in human genomes’, Human Molecular Genetics, 19, R131–R36.

Muyas, F., et al. (2024), ‘De novo detection of somatic mutations in high-throughput single-cell profiling data sets’, Nature Biotechnology, 42 (5), 758–+.

Navin, N., et al. (2011), ‘Tumour evolution inferred by single-cell sequencing’, Nature, 472 (7341), 90–U119.

Nik-Zainal, S., et al. (2016), ‘Landscape of somatic mutations in 560 breast cancer whole-genome sequences’, Nature, 534 (7605), 47–+.

Picardi, E., et al. (2017), ‘REDIportal: a comprehensive database of A-to-I RNA editing events in humans’, Nucleic Acids Research, 45 (D1), D750–D57.

Pogrebniak, K. L. and Curtis, C. (2018), ‘Harnessing Tumor Evolution to Circumvent Resistance’, Trends in Genetics, 34 (8), 639–51.

Sinkala, M. (2023), ‘Mutational landscape of cancer-driver genes across human cancers’, Scientific Reports, 13 (1).

Smit, A. F. A.; Hubley, R.; Green, P. (2013–2015), ‘RepeatMasker Open-4.0’.

Stoler, N. and Nekrutenko, A. (2021), ‘Sequencing error profiles of Illumina sequencing instruments’, NAR Genom Bioinform, 3 (1), lqab019.

Sudlow, C., et al. (2015), ‘UK Biobank: An Open Access Resource for Identifying the Causes of a Wide Range of Complex Diseases of Middle and Old Age’, Plos Medicine, 12 (3).

Sudmant, P. H., et al. (2015), ‘An integrated map of structural variation in 2,504 human genomes’, Nature, 526 (7571), 75–+.

Sun, Z. F., et al. (2017), ‘Indel detection from RNA-seq data: tool evaluation and strategies for accurate detection of actionable mutations’, Briefings in Bioinformatics, 18 (6), 973–83.

Taliun, D., et al. (2021), ‘Sequencing of 53,831 diverse genomes from the NHLBI TOPMed Program’, Nature, 590 (7845).

Tan, M. H., et al. (2017), ‘Dynamic landscape and regulation of RNA editing in mammals’, Nature, 550 (7675), 249–+.

Van der Auwera, Geraldine A., O’Connor, Brian D., and Safari, an O’Reilly Media Company (2020), Genomics in the cloud: using Docker, GATK, and WDL in Terra (First edition. edn.; Sebastopol, CA: O’Reilly Media).

Vázquez-García, I., et al. (2022), ‘Ovarian cancer mutational processes drive site-specific immune evasion’, Nature, 612 (7941), 778–+.

Wiens, M., et al. (2024), ‘Benchmarking bulk and single-cell variant-calling approaches on Chromium scRNA-seq and scATAC-seq libraries’, Genome Research, 34 (8), 1196–210.

Yan, Y. H., et al. (2021), ‘Confirming putative variants at </= 5% allele frequency using allele enrichment and Sanger sequencing’, Sci Rep, 11 (1), 11640.

Yang, L. X., et al. (2013), ‘Diverse Mechanisms of Somatic Structural Variations in Human Cancer Genomes’, Cell, 153 (4), 919–29.

Zafar, H., et al. (2016), ‘Monovar: single-nucleotide variant detection in single cells’, Nature Methods, 13 (6), 505–+.

